# Pooled scanning of protein variants identifies novel RNA-binding mutants

**DOI:** 10.1101/2025.04.02.646914

**Authors:** Julia A. Tasca, John F. Doherty, Emily J. Shields, Shachinthaka Dissanayaka Mudiyanselage, Lauren N. Reich, Kavitha Sarma, Benjamin A. Garcia, Roberto Bonasio

## Abstract

Binding to RNA has been observed for an ever-increasing number of proteins, which often have other functions. The contributions of RNA binding to protein function are best discerned by studying separation-of-function mutants that hamper interaction with RNA without affecting other aspects of protein function. To design these mutants, we need precise knowledge of the residues that contribute to the affinity of the protein for its RNA ligands. Here, we present RBR-scan: a technology to simultaneously measure RNA-binding affinity of a large number of protein variants. We fused individual variants with unique peptide barcodes optimized for detection by mass spectrometry (MS), purified protein pools from single bacterial culture, and assayed proteins in parallel for RNA binding. Mutations in the MS2 coat protein known to impair RNA-binding were correctly identified, as well as a previously unreported mutant, which we validated with orthogonal biochemical methods. We used RBR-scan to discover novel RNA-binding mutants in the cancer-associated splicing regulator SRSF2. Together, our results demonstrate that RBR-scan is a powerful and scalable platform for linking RNA-binding affinity to protein sequence, offering a novel strategy to decode the functional consequences of protein–RNA interactions.

## INTRODUCTION

RNA-binding proteins (RBPs) dynamically associate with RNA to form ribonucleoprotein (RNP) complexes that sustain cellular homeostasis^1^. Due to their extensive involvement in biological processes, mutations in RBPs can cause cancer, neurological disorders, and other major diseases^2^. The distinct molecular mechanisms that govern protein–RNA interactions are incompletely understood, but numerous biochemical and functional studies have provided important insight into RBP structure^2–4^. RBPs were initially classified by the presence of one or more known RNA-binding domains (RBDs), such as the RNA recognition motif (RRM), K-homology (KH) domain, or double-stranded RNA-binding domain^5,6^. However, major strides in RNA interactome capture approaches, many of which utilized ultraviolet (UV) crosslinking-mass spectrometry (XL-MS), have uncovered thousands of novel RBPs that lack canonical RBDs^7–14^.

In addition to identifying new RBPs, some of these technologies, including our own RNA-binding region identification (RBR-ID)^10^ approach, provided “mapping” information, typically reporting on the position of the protein–RNA crosslink at domain, peptide, or individual amino acid resolution^7–14^. Protein–RNA crosslinks marked potential interaction sites, which guided the design of targeted deletion mutants that provided insights into the biological and biochemical functions of these interactions^15–17^. Thus, XL-MS–based methods have advanced our understanding of RBP structure and function, though these approaches are not without limitations.

UV crosslinking introduces a covalent bond between protein and proximal RNA at “zero distance”, preserving protein–RNA interactions under denaturing conditions. However, it introduces bias toward amino acids with high photoreactivity^5,6,14,18^. Residues with side chains containing sulfur, aromatic rings, or π-bonds (e.g. Cys, Tyr, Trp, Phe, Met, His) have the highest crosslinking efficiency, while others with low photoreactivity, like aliphatic residues (e.g., Leu and Ile) are underrepresented despite their role in RNA binding^14,18^. For example, aliphatic residues contribute to protein–RNA hydrogen bonding in KH domains, yet crosslink poorly^3,4,19^. Consequently, UV crosslinking has a limited capacity for identifying functional residues with low photoreactivity.

Another limitation of XL-MS methods is incomplete protein sequence coverage. Low protein abundance and poor MS detectability of peptides resulting from suboptimal peptide length, charge, or hydrophobicity contribute to this challenge^20–25^. The median coverage of a human protein from a trypsin-digested proteome is 56%, increasing to only 79.2% even when trypsin is combined with five additional proteases^22^. Moreover, ∼20% of RBPs interact with RNA through arginine- and lysine-rich intrinsically disordered regions (IDRs)^26^, which standard, bottom-up mass spectrometry cannot detect. Since trypsin cleaves at every arginine and lysine, many critical RNA-binding residues within these regions remain undetected. As mass spectrometry remains the predominant high-throughput method for characterizing RBPs, its inherent limitations pose a major barrier to comprehensive mapping of RNA-binding interfaces.

To meet the demand for high-sensitivity functional mapping of protein–RNA interactions, we created RBR-scan: a technology that scans proteins for functional RNA-binding residues using genetically coupled peptide barcodes. By purifying a pool of mutants from a single bacterial culture, we were able to assess defects in RNA binding in a large number protein variants at the same time. RBR-scan improves upon previous methods by overcoming challenges such as detecting residues with low photoreactivity and increasing coverage of binding sites that may be missed by other technologies. Since the peptide barcodes used in RBR-scan are optimized for liquid chromatography-mass spectrometry (LC-MS), they bypass issues of poor detection and low coverage of certain peptides from proteins that are challenging for traditional MS methods. Unlike crosslinking-based approaches, RBR-scan tests the affinity of a protein for its RNA ligand in native conditions. This method not only allows for a more accurate representation of the protein–RNA interaction landscape, but also preserves the true electrostatic properties of the binding sites, providing a more authentic view of how these interactions occur. We applied RBR-scan to map functional RNA-binding residues within MS2 coat protein and SRSF2, revealing new insights into the RNA interaction landscapes of these proteins.

## RESULTS

### Peptide barcodes can serve as a readout for RNA binding

Based on previous studies that used peptide barcodes for directed evolution of nanobodies^27^, we posited that peptide barcodes could also be used to distinguish defects in RNA binding within pool of protein variants (**Fig. 1A**). To explore this possibility, we used the well-characterized MS2 bacteriophage coat protein and its cognate stem loop RNA as a model^28–31^. We selected three mutants previously described to impair RNA binding to MS2, including two mutants with single point mutations (N55D or K61A) and one mutant with three point mutations in the RNA–binding pocket (R49A-K57A-K61A, “3A”)^28^. DNA fragments encoding these three MS2 coat protein mutants as well as the wild type (WT) sequence each fused with a distinct N-terminal peptide barcode (**Table S1**) were commercially synthesized (**Fig. 1B**). The variants were individually cloned, expressed, and purified from *E. coli* cultures (**Fig. 1C, S1A**). Barcode-mutation pairs from individual plasmids were verified by Sanger sequencing. We also validated barcode-mutation pairs from a pool of the four plasmids via Oxford nanopore sequencing, a long-read sequencing platform providing single-read coverage of both the peptide barcode and the corresponding mutant codon (**Fig. S1B–C**, **Table S1**). The pairing of each mutant MS2 coding sequence with the expected peptide barcode was nearly perfect (**Fig. S1B**), with a very small percentage of reads showing unexpected sequences, likely due to the error rate in nanopore sequencing (**Fig. S1C**). To verify the mutant phenotypes, biotinylated MS2 stem loop (MS2 SL) RNA was synthesized *in vitro* and incubated with the four individual MS2 variants, and biotinylated RNA– protein complexes were purified with streptavidin (**Fig. 1D, Fig. S1A**). Western blot analysis of individual pull-downs confirmed the severe defects in RNA binding of the N55D, K61A, and R49A-K57A-K61A MS2 mutants (**Fig. 1D**).

**Figure 1.**
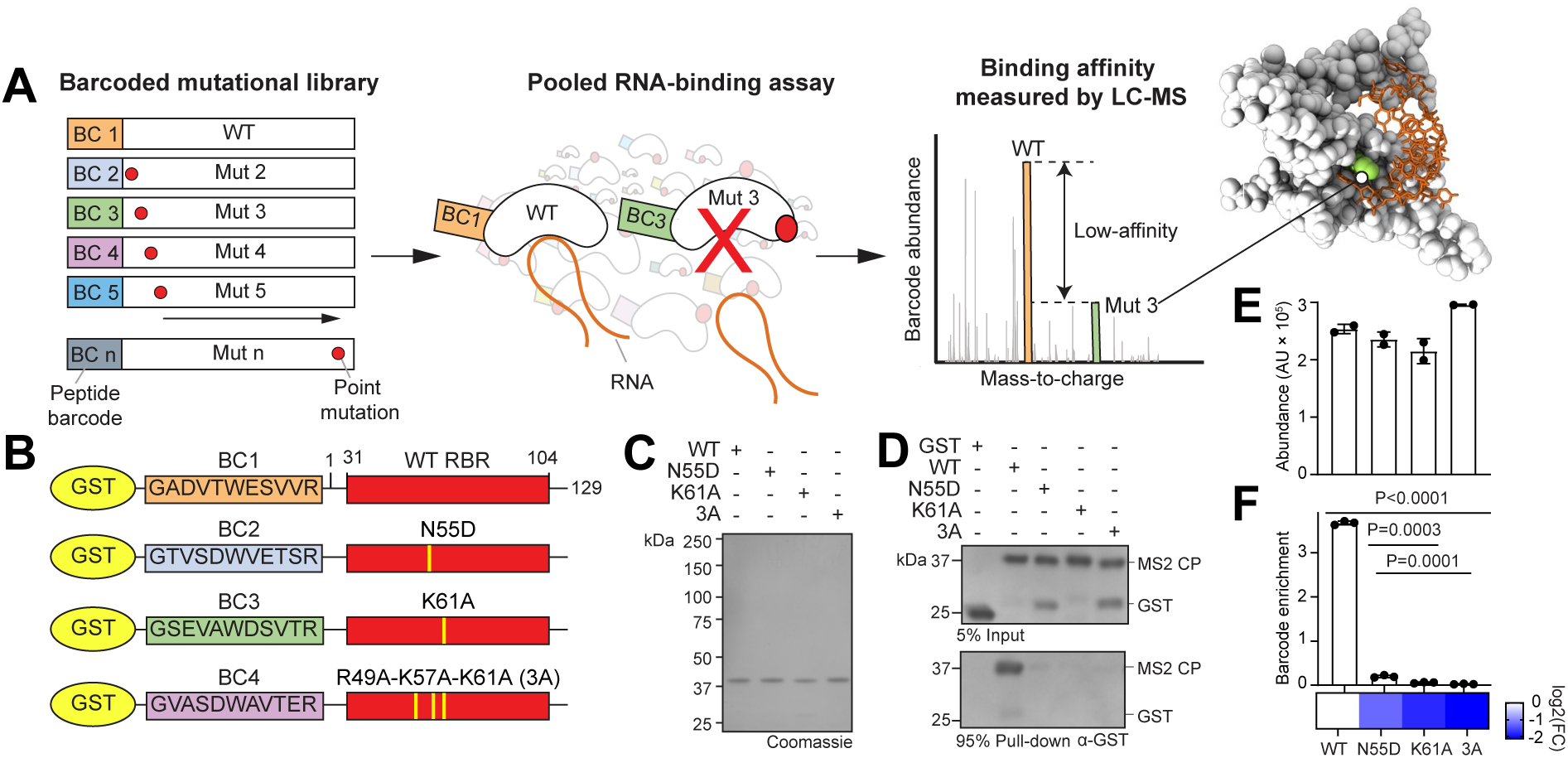
Peptide barcodes allow testing of multiple protein variants in a single assay. (A) Experimental scheme of RBR-scan. RBR-scan plasmid libraries comprise multiple genes with point mutations fused to distinct peptide barcodes. Pools of protein variants are purified and subjected to RNA-affinity purification. The abundance of peptide barcodes in the RNA-bound vs. the input fraction is determined by LC-MS. Mutants with RNA-binding defects will show depletion of the corresponding barcode in the RNA-bound fraction relative to WT barcodes. The structure of the MS2 coat protein (PDB: 1ZDI) and its cognate stem loop RNA (orange) are shown with a key RNA-binding residue highlighted in green. (B) Domain organizations of GST-fused MS2 coat protein variants with N-terminal peptide barcodes. See methods for barcode composition rules. (C) Coomassie stain of GST-fused barcoded MS2 coat proteins (predicted MW: 43.5 kDa). (D) Western blot of individual MS2 coat protein variants after RNA affinity purification. (E) Relative peptide barcode abundances as measured by LC-MS in an equimolar pool of the four protein variants. Bars represent the mean ± SEM. Data from 2 technical replicates. (F) Same as (E) but after RNA affinity purification. Bars represent the mean barcode enrichment ratio (i.e., ratio of input to pull-down) relative to input ± SEM. Heatmap shows the log_2_(fold-change) of the enrichment ratio relative to WT. Data from 3 technical replicates. P values are from ANOVA and Holm-Sidak test.

To determine whether we could measure binding defects within a pool of mutants using peptide barcodes, to simulate the high-throughput approach, we combined the barcoded MS2 coat protein variants in an equimolar protein pool and performed the pull-down as described above. We then analyzed both input and RNA-bound fraction using LC-MS. The four barcodes, and therefore the 4 MS2 coat protein variants, were present in comparable amounts in the input (**Fig. 1E**), whereas barcodes associated with the three RNA-binding mutants were strongly depleted relative to WT in the RNA-bound fraction (P<0.0001) (**Fig. 1F**). We also detected significant differences in abundance between N55D and the two other RNA binding mutants (K61A, P=0.0003; R49A-K57A-K61A, P=0.0001). Based on these results, we concluded that barcode quantification was effective in reporting RNA binding defects of specific variants within a protein pool.

These findings collectively indicate that quantification of peptide barcodes after RNA-based affinity purification from protein variant pools can reliably identify mutant proteins defective for RNA binding.

### Design of barcoded mutant pools

Having demonstrated that RNA affinity purification from a mixture of four protein variants successfully identifies mutants with RNA-binding defects, we next sought to generate a much larger pool interrogating several known and uncharacterized mutants simultaneously.

To generate these libraries, we first designed a set of peptide barcodes with optimal properties for LC-MS detection, as previously described^27^, but with some modifications. Each barcode was composed of 11 amino acids, designed to maximize LC-MS detectability by considering factors such as solubility, a +2 charge, and a hydrophobicity index ranging from 5 to 45, based on previously defined parameters^27,32^ (**Fig. S2A**). Within these constraints, there were 5.4 × 10^11^ theoretical peptide sequences available for barcoding. Given the extent of theoretical barcode diversity, we limited the upper threshold of molecular weight to 1.5 kDa (similar to FLAG or 6xHis) to minimize variations in purification or effects on function^33–35^.

Based on these parameters, we designed 155 peptide barcodes *in silico* to be tested for their detection efficiency by LC-MS (**Fig. S2B; Table S2**). We fused all 155 barcodes to the same WT MS2 coat protein sequence by PCR, then cloned the pool in an expression vector and purified from a single *E. coli* culture. The relative abundance of the barcodes at the DNA level was quantified with next-generation sequencing to correct for possible biases during the cloning process. We then measured the abundance of the barcodes at the protein level. Because all barcodes were fused to the same protein sequence, we reasoned that deviation from a linear correlation between DNA abundance and peptide abundance would indicate unexpected variations in detection efficiency that might negatively impact our downstream RBR-scan assay. Therefore, we discarded 53 barcodes with the largest deviation from linearity and retained the remaining 102 barcodes for subsequent RBR-scan experiments (see methods) (**Fig. S2C**).

Because of the increased complexity of these libraries, we implemented an existing strategy^27^ that would facilitate the separation of the barcodes from the rest of the protein after purification (**Fig. S2D**). Each variant in an RBR-scan pool is fused at the N terminus with: 1) an MBP affinity tag for purification of the pool after expression in *E. coli*, 2) a distinct peptide barcode sequence optimized for mass spectrometry (see above), and 3) a 6xHis tag for peptide barcode enrichment. Two PreScission protease (PSP) sites enable proteolytic separation of the barcode from the MBP moiety and from the protein bait (**Fig. S2D**). Additionally, the arginine residue in PSP site #1 and the invariant arginine at the C terminus of each barcode permits for the isolation of the peptide barcodes by trypsin cleavage before LC-MS analysis (**Fig. S2A, S2D**). This also allows for release of the barcode peptides by on-bead trypsinization making the 6xHis-enrichment and PSP cleavage steps optional. Each workflow offers distinct advantages: barcode enrichment enhances sensitivity, while on-bead trypsinization provides a faster, streamlined protocol.

### High-throughput generation of barcoded protein libraries

We invented RBR-scan to investigate a large number of protein variants simultaneously to identify residues within an RBP that contribute to RNA binding with minimal or no pre-existing information on their potential position. At this scale, synthesis of each individual DNA fragment as described above is very costly with existing technologies; therefore, we devised a method to generate all desired mutants paired with distinct individual barcodes using a modified overlap-extension (OE-) PCR that minimized cost and labor (**Fig. S2E, Table S3**).

Traditional mutagenesis by OE-PCR^36^ introduces a mutation of interest by generating overlapping mutagenized fragments using WT DNA as a template and primers containing the mutation of interest. A necessary but time-consuming step in the conventional method is the removal of the WT template from the 5’ and 3’ fragments by gel purification before combining them to synthesize the full-length mutant gene in a second PCR reaction. This step is required to avoid the amplification of the original WT sequence. We expected that we could bypass gel purification by utilizing a template where all deoxy-thymidines (dTs) had been replaced with deoxy-uridines (dUs), which allows for its selective cleavage via a mixture of uracil DNA glycosylase (UDG) and endonuclease III, commercially available as “uracil-specific excision reagent” (USER)^37,38^ (**Fig. S2E**). PCRs to generate overlapping mutant fragments were conducted in two separate 96-well plates. In one plate, barcoded fragments encoding the N-terminal portion of the protein up to the mutation (“AB” fragments; **Fig. S2E**) were generated using a forward primer containing a constant annealing region to the gene and a peptide barcode overhang (primer “a”) and a reverse primer containing the desired point mutation (primer “b”). DNA fragments encoding the remaining portion of the protein, from the desired mutation to the C-terminus, were generated in a second plate (“CD” fragments from primers “c” and “d”; **Fig. S2E**) using a forward primer encoding the mutation and a reverse shared primer complementary to the 3’ end of the protein-encoding sequence in the template. After the first amplification, we degraded the WT template by USER treatment (**Fig. S2F**), skipped the gel-based purification and directly combined the overlapping DNA fragments along with external primers (“e” and “f”) to undergo OE-PCR and generate full-length barcoded mutants.

We validated the specificity of our PCR strategy by demonstrating that only overlapping fragments could form a full-length product, whereas mismatched fragments could not (**Fig. S2G**). Next, we wanted to confirm that the USER reaction fully degraded the WT template. For this, we introduced a HindIII recognition site in the internal primers encoding the W32A mutation so that only the correct product and not the WT sequence could be digested by HindIII. Upon HindIII restriction enzyme treatment of W32A OE-PCRs with and without the HindIII site, we established that the sole product at the predicted size was the OE-PCR mutant (**Fig. S2H**).

With this high-throughput OE-PCR strategy, we generated a library encoding barcoded MS2 coat proteins with mutations affecting 15 residues with known RNA-binding phenotypes (as controls), as well as 25 residues not previously tested in the context of RNA binding. Each pool contained a total of 96 barcoded genes representing 47 different MS2 coat protein variants (including WT) (**Fig. 2A**). To minimize barcode- and codon-dependent biases in the expression, purification, or detection of the protein variants, each mutation was represented by two distinct peptide barcodes linked to two different codons for the same mutant amino acid (**Table S1**). As a control, we fused the WT sequence to four different barcodes in each pool. To further increase reproducibility and minimize bias, we generated two separate plasmid libraries yielding two independent protein pools comprising the same mutants linked to different barcode combinations. In total across the two pools, 4 different barcodes were linked to each mutant, and 8 barcodes were linked to the WT protein (**Table S1**). Individual OE-PCR reactions were analyzed on an agarose gel (**Fig. S3A**), then pooled and cloned into an expression vector. The resulting plasmid pool contained only plasmid with insert and not empty expression vector, as determined by PCR (**Fig. 2B**).

**Figure 2.**
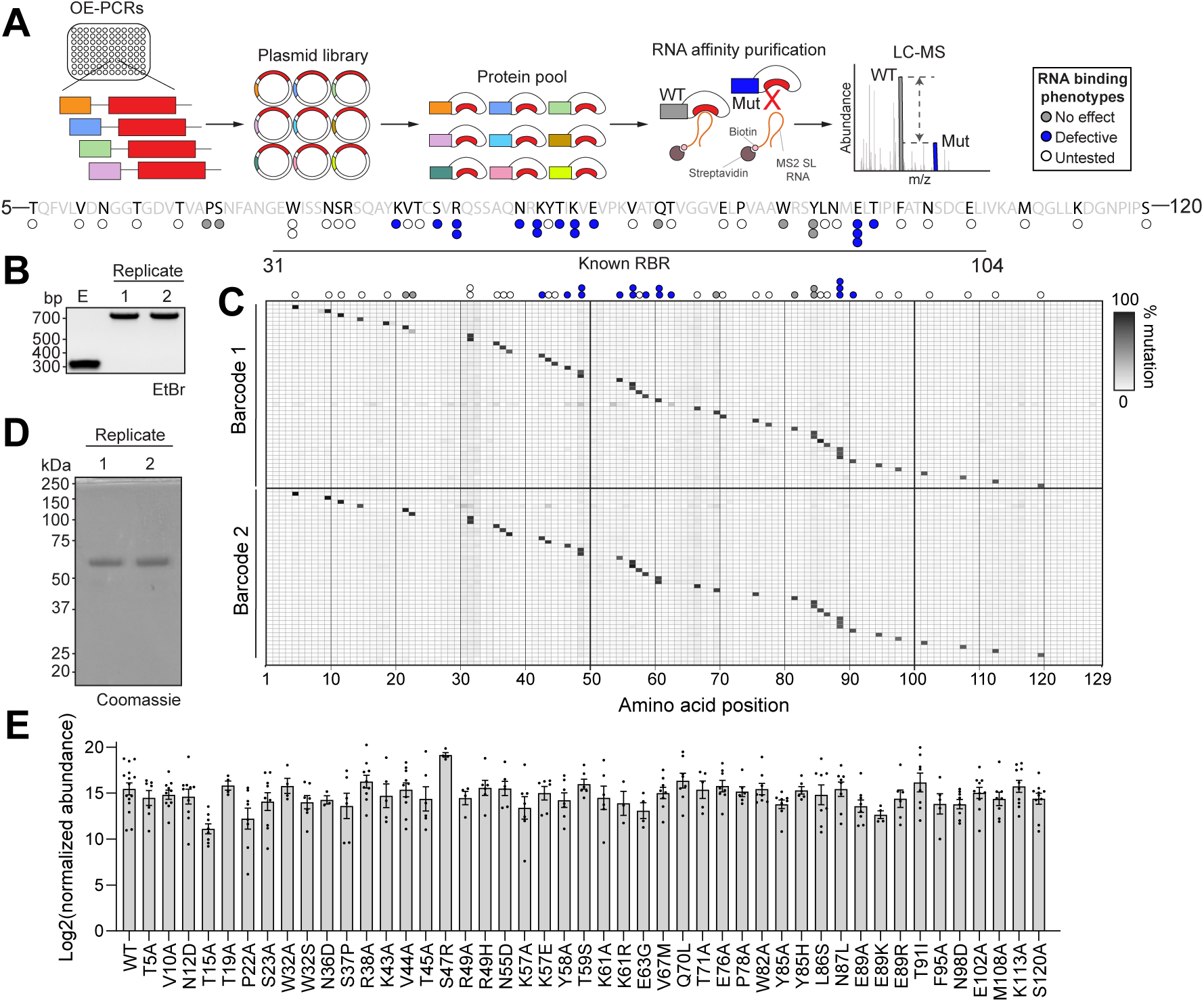
Construction of peptide barcode mutant pools by PCR. (A) Scheme of high-throughput RBR-scan for MS2 coat protein. Peptide barcodes assigned to mutations were fused to the corresponding protein variant by a modified two-step OE-PCR (**Fig. S2E**). Barcoded DNA libraries were pooled and cloned into an expression vector in batch. Protein pools were expressed and purified from a single bacterial culture. Barcoded protein pools were incubated with biotinylated stem-loop RNA, followed by streptavidin-mediated affinity purification of protein–RNA complexes. RNA-binding defects were assessed by LC-MS analysis of peptide barcode abundances relative to WT. Residues selected for RBR-scan analysis are annotated on the MS2 coat protein primary sequence (UniProt P03612). Circles represent location of point mutations for RBR-scan pools. Mutants previously tested that showed no effect on RNA binding are in gray; known RNA binding mutation in blue; untested mutations in white). (B) PCR products from two RBR-scan plasmid libraries (expected size: 754 bp) and an empty vector (expected size: 278). E, empty vector. (C) Mutation frequencies (% of reads) of each amino acid (x-axis) along the MS2 coat protein sequence. Barcoded MS2 coat protein variants are shown (y-axis), and barcodes 1 and 2 denote two different peptide barcodes and codons for each amino acid within the same pool. Positions of intended mutations are indicated at the top of the heatmap, following the same color scheme as Fig. 2A for RNA binding phenotypes. (D) Coomassie stain of MBP-fused peptide barcoded MS2 coat protein pools (predicted MW: 60.5 kDa). (E) Mass spectrometry analysis of peptide barcode abundances MS2-coat protein pools. Point mutations indicated on x-axis. Bars represent the mean log_2_(barcode abundance). Dots represent abundances from individual barcodes across replicates. Data from 4 biological replicates.

The effective pairing of specific barcodes to each variant in the plasmid pools was verified using nanopore sequencing (**Fig. 2C, S3B–C**). Out of a total of 96 barcode configurations, all 47 protein variants (including WT) were represented in the DNA libraries (**Fig. 2C, S3B**). The vast majority of barcodes present in the pool (90/96 in barcode mutation set 1 and 92/96 in barcode mutation set 2) were paired with the correct amino acid change (see methods). For each barcode, a fraction of the reads (ranging from 5% to 45%, but in most cases < 25%) displayed WT sequence (**Fig. S3C**). Since degradation of the template by USER was complete (**Fig. S2F–H**), we suspect that WT sequences paired with barcodes for mutant variants arose by recombination during PCR or cloning^39^. As shown below, this small amount of WT sequence contaminants did not impede our ability to identify loss-of-function mutations. An even smaller proportion of reads contained unexpected mutations, very likely due to sequencing errors by nanopore (as also seen for the synthetic genes in **Fig. S1C**). Barcode-mutation combination not detected by nanopore were not further considered during the LC-MS analysis.

RBR-scan protein pools were expressed and purified from single *E. coli* cultures (**Fig. 2D**) and analyzed by LC-MS. Peptide barcode analysis showed that all 47 variants (including WT) were present in the protein pools (**Fig. 2E**), although some individual barcodes were missing. Missing and low abundance barcodes correlated well with our result from nanopore, confirming that none of the selected barcodes negatively affect protein expression and purification (see **Fig. S2**). Peptide barcodes for all 47 variants were present in the input (**Fig. 2E**). Our results demonstrate that peptide barcodes can be genetically linked to specific mutations in a high-throughput fashion with minimal cost and labor using a modified OE-PCR.

### Identification of multiple known RNA-binding mutants in parallel

To determine if peptide barcode abundance from mutant pools generated by OE-PCR correlated with expected phenotype (**Table S4**), RBR-scan protein pools were affinity purified with MS2 SL RNA and analyzed by LC-MS (**Fig. 2A, 3A**). We predicted that barcodes from mutants known to impact RNA binding would be depleted, whereas barcodes from WT-like controls would not. As expected, barcodes from WT-like controls such as S37P, V67M, and Q70L were not depleted upon RNA pull-down. On the other hand, barcodes from 14 of the 17 known RNA-binding mutants were depleted, including N55D and K61A (**Fig. 1F, 3A** and **Table S4**)^28^. These results demonstrate that changes in peptide barcode abundance after RNA pull-down of RBR-scan protein pools can identify protein variants defective in RNA binding.

### RBR-scan identifies a novel MS2 coat protein mutation

While RBR-scan correctly identified mutations with known RNA-binding defects, such as N55D, K61A, R49A, and N87L, some barcodes linked to substitutions at residues not previously known to affect RNA binding were also consistently depleted. For example, W32, a residue at the N-terminal edge of the annotated RBR (**Fig. 2A**), exhibited binding defects by RBR-scan when substituted with two different amino acids (W32A and W32S, **Fig. 3A**). To evaluate the loss-of-function of W32A, we expressed and purified recombinant MS2 coat protein carrying this point mutation (**Fig. S4A**) and analyzed its interaction with cognate RNA by RNA-mediated pull-down followed by western blot (**Fig. 3C**). In agreement with the observation made with RBR-scan, the W32A failed to bind to the MS2 RNA, to the same extent as the known RNA-binding defective mutant N55D (**Fig. 1D**). To further validate our results from the pull-down, we assessed protein– RNA complex formation by electrophoretic mobility shift assays (EMSAs) (**Fig. 3C–D, S4B**) and quantified binding by fluorescence polarization (FP) (**Fig. 3E–F**). WT MS2 coat protein exhibited binding at nanomolar affinity for MS2 SL RNA, whereas neither N55D nor W32A displayed any detectable affinity for the RNA. Confirmation of known RNA-binding phenotypes along with the discovery of a novel mutant underlines the utility of RBR-scan as a tool to scan proteins for residues that contribute to their affinity for RNA.

**Figure 3.**
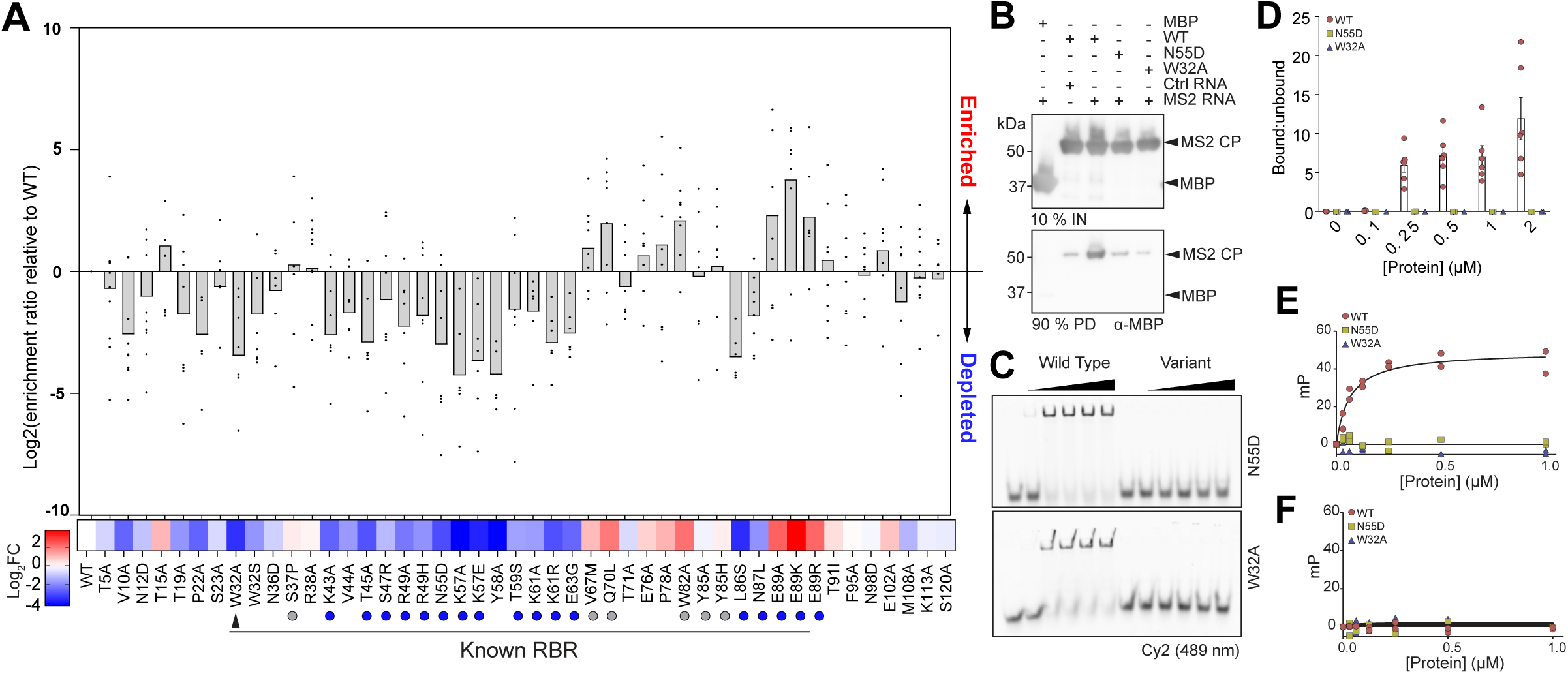
Identification of a previously unknown RNA-binding mutant in the MS2 coat protein. (A) Mass spectrometry analysis of peptide barcodes after RNA pull-down. Mutations are listed on the x-axis. Known RNA-binding point mutations (red circles), WT/WT-like mutations (grey circles), and putative RNA-binding mutant (black triangle) annotated. Bars and heatmap represent the mean enrichment ratio relative to WT. Dots represent enrichment ratios of individual barcodes across replicates. Data from 4 biological replicates. (B) Western blot of individual MS2 coat protein mutants after RNA pull-down with MS2 stem loop RNA or control RNA. (C) EMSAs with titrations of recombinant MS2 coat proteins and 50 nmol Alexa488-MS2 stem loop RNA in the presence of 1 g/L yeast tRNA. Complexes were separated on native gels, and RNA was imaged by Cy2 excitation. Data is representative of ≥2 experiments. (D) Densitometric quantification of the proportion of RNA bound in (C). (E–F) Fluorescence polarization assays of individual MS2 coat protein mutants with MS2 stem loop RNA (E) or control RNA (F). Data from 2 technical replicates.

### RBR-scan identified a novel RNA-binding mutant in SRSF2

Having validated our approach by identifying a novel RNA-binding mutant in the MS2 coat protein, we next sought to investigate functionally important RNA-binding residues in the context of human disease. We selected SRSF2, a known RBP and core spliceosomal component, whose dysfunction is implicated in cancer. SRSF2 binds exonic splicing enhancer sequences on pre-mRNA (**Fig. 4A**)^40^, and its misregulation is linked to various diseases including chronic myeloid leukemia^41^, which involves mutations at a critical RNA binding residue, proline 95. Structurally, SRSF2 contains an RRM and a C-terminal RS domain, thought to mediate RNA binding and protein–protein interactions, respectively^42^.

**Figure 4.**
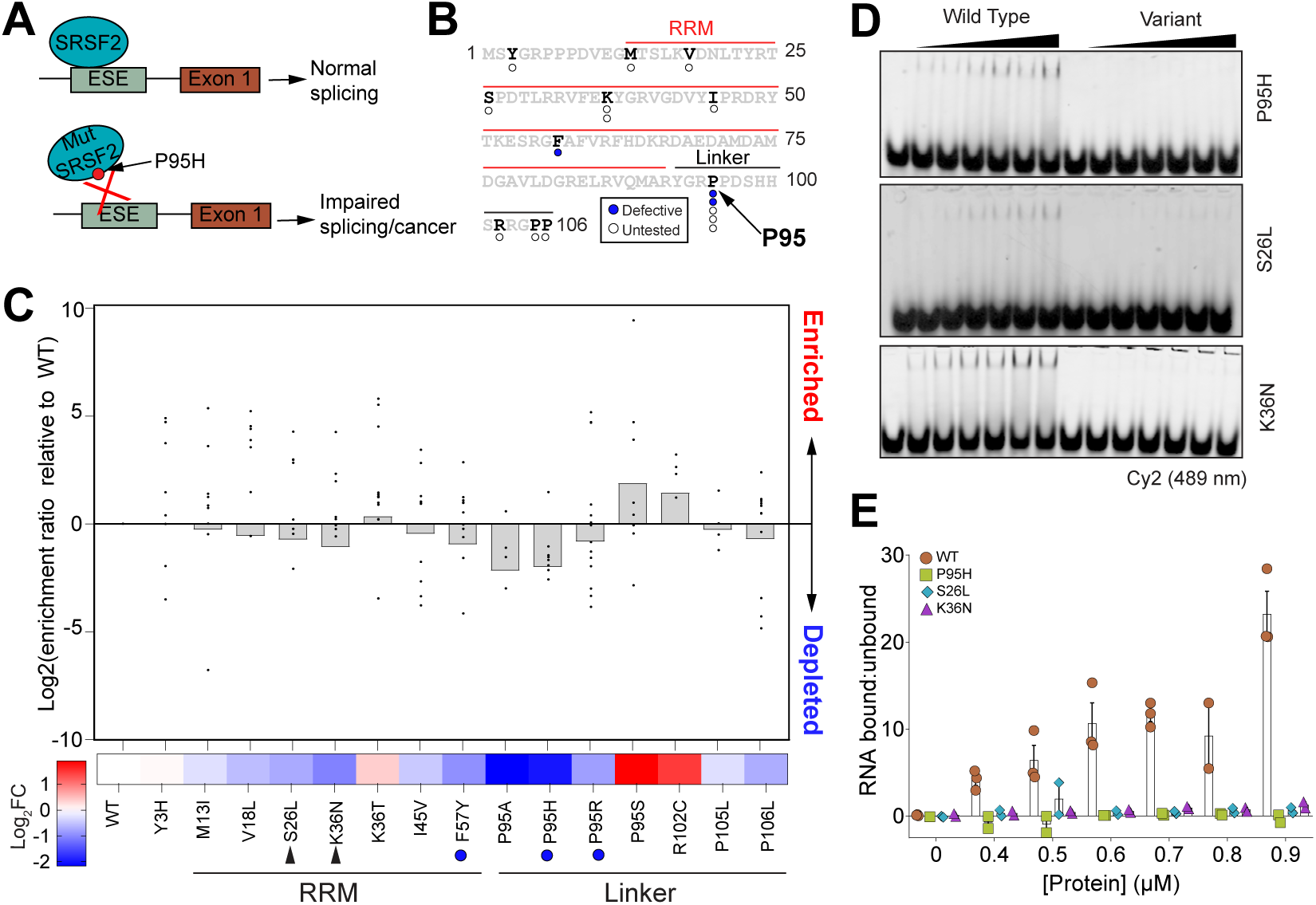
Identification and validation of RNA-binding mutations within SRSF2. (A) Schematic for SRSF2 function in splicing with and without P95H mutation. Mut, mutant. ESE, exonic splicing enhancer. (B) SRSF2 residues selected for RBR-scan are annotated on the protein primary sequence along with SRSF2 domain organization (UniProt Q01130). Circles represent location and expected phenotypes of point mutations for RBR-scan pools (white, untested; blue, defective for RNA binding). (C) Mass spectrometry analysis of peptide barcodes from RNA-affinity purified proteins. All RBR-scan mutations are listed on the x-axis. SRSF2 domains (UniProt Q01130), known RNA-binding point mutations (red circles), WT (grey circle), and unreported RNA-binding mutants for individual analysis (black triangle) are annotated. Bars and heatmap represent the mean enrichment ratio relative to WT. Dots represent enrichment ratios of individual barcodes across replicates. Data from 8 biological replicates. (D) EMSAs with titrations of recombinant SRSF2 RRMs and 50 nmol Alexa488-MELK RNA in the presence of 1 g/L yeast tRNA. Complexes were separated on native gels, and RNA was imaged by Cy2 excitation. Data is representative of ≥2 experiments. (E) Densitometry-based quantification of the ratio of bound:unbound RNA from (D). Bars show means ± SEM from ≥2 experiments.

To identify loss-of-function mutations in SRSF2, we introduced 16 point mutations within a defined region of SRSF2 (UniProt Q01130), including the RRM and linker (**Fig. 4B**). Mutations were selected if they exhibited known or predicted pathogenic consequences based on UniProt’s genetic variant database^43^. The RNA bait was comprised of 3 repeats of an SRSF2-binding motif (UCCAG) from *MELK* RNA, a known target of SRSF2^44^. The full-length SRSF2 RBR-scan pools contained 32 barcoded variants with two different barcode configurations resulting in 4 distinct barcodes per mutant and 8 possible WT barcodes (**Table S1**). Barcode-mutation combinations in the plasmid pool were verified by nanopore sequencing (**Fig. S5A**), and undetected barcodes were excluded from further analyses. We also confirmed that the RBR-scan plasmid pool contained only plasmid with SRSF2 insert by PCR (**Fig. S5B**).

Protein purity was assessed by Coomassie stain (**Fig. S5C**) and confirmed that all 16 variants and WT were detected by LC-MS in the input fraction (**Fig. S5D**). We observed depletion relative to WT after RNA affinity purification of known RNA-binding mutants including F57Y (part the conserved RNP1 motif^42,45^), P95H^44^, and P95R^46^ (**Fig. 4C**).

Barcodes associated with residues in the RRM not previously reported to affect RNA binding, such as S26 and K36, were also depleted (**Fig. 4C**). To validate these two putative novel RNA-binding residues identified within the RRM, we expressed and purified recombinant SRSF2 RRMs (truncated before the RS domain) carrying S26L or K36N point mutations, as well as WT and P95H RRMs as negative and positive controls, respectively (**Fig. S5E**). RNA-binding affinities were assessed by EMSA with a single fluorescently labeled repeat of the MELK motif (**Fig. 4D– E**, **S5F**, **Table S4**). Both S26L and K36N exhibited a decrease in binding affinity relative to WT. Our results demonstrate that RBR-scan correctly identified known and novel RNA-binding mutations in a human splicing protein linked to cancer.

## DISCUSSION

A crucial step toward understanding RBP function is to identify the amino acid residues that contribute to RNA–binding affinity. To preserve protein structure while exploring RNA binding mechanisms, separation-of-function mutants with the most minimal effective mutation are required. Mutational libraries of peptide barcoded-RBPs allowed us to scan for residues that contribute to RNA binding along the full length of a protein. The MS2 coat protein was used to demonstrate proof-of-concept for RBR-scan because it is a well-studied protein with 3D structures available to visualize protein–RNA interactions ^28,30,31^. Despite the extensive research on protein– RNA interactions in the MS2 coat protein, RBR-scan identified a novel mutation (**Fig. 3A**) that diminished RNA binding (W32A, **Fig. 3B–F**, **S4B**). RBR-scan also detected known and novel mutations in the human splicing factor SRSF2 (S26L and K36N, **Fig. 4D–E, S5G**). Collectively, these results emphasize the capabilities of RBR-scan as a technology that can be used to discover novel protein–RNA interactions.

### RBR-scan fills a technological gap

Protein–RNA crosslinking methods have uncovered many new RBPs and, in some cases, helped design deletion mutants to investigate the functional consequence of RNA interactions^15^. However, such methods do not provide functional information about protein–RNA interactions and are limited by low resolution and bias. In addition, mutant design guided by RNA interactome capture involves large deletions or extensive trial-and-error to narrow down the most minimal effective mutation without necessarily obtaining a complete separation-of-function phenotype^15^. In this scenario, expression, purification, and testing of individual proteins is required, which cannot be easily scaled to test tens or hundreds of separation-of-function mutants.

Existing methods^27,47,48^ have used peptide barcoding for multiplexed functional characterization of protein variants. The most notable of these methods is NestLink^27^, which was developed for deep mutational scanning of synthetic nanobodies to identify mutants with improved binding characteristics. NestLink pioneered crucial experiments such as LC-MS-optimized peptide barcode design and high-throughput functional screening. However, constraints in library design make NestLink incompatible with RBP functional analyses. For example, NestLink randomly combines mutant and barcode libraries that must be sequenced to determine mutant-barcode pairings. Their workflow is restricted to a maximum of 600 cycles making it impossible to adapt this method to a protein longer than 200 amino acids, which is much smaller than a typical human protein^49^. Due to the massive diversity of their barcode library (∼100 million), only a very small fraction of barcodes were validated experimentally. Here, we show that the performance of *in silico* optimized barcodes varies greatly (**Fig. S2B, S2D**), even when parameters such as hydrophobicity, charge, and size are kept within predetermined ranges, underlining the importance of empirical validation of peptide barcodes prior to library design.

### Advantages and limitations of RBR-scan

We created RBR-scan to address the need for a high-throughput, unbiased technology to scan RBPs of any length for mutants that affect RNA binding. Using a peptide barcoding strategy, we purified and assayed close to fifty protein variants from a single bacterial culture (**Fig. 2D–E**). This approach could be easily scaled up to test hundreds of mutations simultaneously, given that routine mass spectrometry analysis of complex proteomic samples can yield tens of thousands of unique peptides within a wide dynamic range spanning 10 orders of magnitude ^50,51^.

RBR-scan also addresses the shortcomings UV crosslinking methods. RNA affinity purification of barcoded protein libraries under native conditions constitutes a more direct method to test protein variants for their affinity to RNA under native conditions. Consequently, RBR-scan can identify contributions to RNA binding by residues that do not crosslink well^4^, are within protein regions difficult to detect by MS, or do not directly contact RNA^52,53^. RBR-scan should not be regarded as a replacement, but rather a complement to existing methods. RNA interactome capture methods typically cannot identify specific residues involved in protein–RNA interactions, but they can identify putative RBPs in a high-throughput manner. These candidate RBPs can then be scanned to narrow down the possible residues that affect binding affinity, substantially reducing the number of separation-of-function mutants to validate in biological assays.

PCR mutagenesis has been widely used to introduce substitutions, insertions, and deletions into genes. Traditionally, PCR mutagenesis involves synthesis of complementary fragments with the mutation of interest from a WT gene template. Isolation of these fragments from the WT template is crucial but requires a time-consuming gel purification step that cannot be easily scaled up. RBR-scan drastically reduces the amount of time it takes to synthesize a mutational library by removing the gel purification step (**Fig. S2E**). We showed that by replacing all of the dTs with dUs in the WT MS2 coat protein template, we could enzymatically degrade the template (**Fig. S2F–H**) and generate 96 distinct barcode-variant pairings in 3–4 hours. The unique peptide barcodes added to the 5’ end allowed us to pool all 96 sequences after the OE-PCR, eliminating the need to handle individual clones during all subsequent steps. By coupling RBR-scan with nanopore sequencing, we can verify barcode-mutation linkages within the same read, without constraints on gene size—an improvement over existing methods limited by shorter read lengths.

We note that nanopore sequencing is an evolving technology, and sequencing error occurs at a higher rate than in Illumina sequencing^54^. However, we expect that higher accuracy nanopore analysis pipelines will further improve our analyses in the future^55^. As shown in **Figures 3B** and **4C**, RNA binding mutants exhibited the expected phenotypes after RNA-affinity purification, suggesting that the small contamination of WT reads in the barcoded pools does not meaningfully affect our assay.

RBR-scan was designed to probe the function of one amino acid at a time, as a high-throughput modern version of alanine scanning^56^. In some cases, individual substitutions are not enough to substantially reduce binding. For instance, E89 and N87 MS2 coat protein residues regulate RNA–binding specificity and affinity, and mutational studies showed that N87S-E89T decreases binding affinity more than the individual mutants alone^57^. In a broader sense, coordination of multiple near-by residues within known RBDs facilitate RNA–binding recognition of RNA^4,58^. In addition, IDR-containing proteins undergo conformational changes upon RNA–binding which may bring together sequences that are distant in the primary structure^59^. Thus, RBR-scan is best used to assess many point mutants simultaneously, a subset of which can then be combined for further testing. In fact, with appropriate primer design, it would be straightforward to design RBR-scan libraries to interrogate combination of mutants, provided that the residues are close enough to be spanned by a single PCR primer (i.e. within ∼100 nucleotides, based on current technologies^60^).

### Outlook and conclusion

RBR-scan is a powerful new technology that allows for simultaneous testing of hundreds of protein variants. The mutants can span entire proteins without limitations in length but can also be focused on a specific protein region with saturation mutagenesis, for gain-of-function studies. While demonstrated here for RNA binding mutants, RBR-scan can be used to investigate the effect of mutations on any protein property that allows for the physical separation of functional vs. non-functional variants, both *in vitro* and also *in vivo*.

## EXPERIMENTAL PROCEDURES

### Plasmids and sequences

The construction of the pGEX-6P1 backbone was described previously^61^. The DNA sequences for MS2-CP variants (WT, N55D, K61A, K49A-K57A-K61A) were synthesized by Azenta and subcloned into the pGEX-6P1 expression vector. pMAL-c5P was based on pMAL-c5X (NEB, N8114), which contains a PreScission Protease cleavage site instead of factor Xa. Truncations and individual mutants were obtained by PCR. Full length human SRSF2 was codon optimized for *E. coli* and synthesized by IDT. All oligonucleotide and synthetic DNA sequences can be found in **Table S3**.

### Antibodies

The following antibodies were used for western blot: GST (Cell Signaling Technology, 3369S) and MBP (NEB, E8032S).

### RBR-scan design

Each RBR-scan variant contains an N-terminal MBP affinity tag, a modified N-terminal PreScission Protease site (PSP #1, LEVLFQGP<u>R</u>), a unique peptide barcode, a His tag, a C-terminal PreScission Protease site (PSP #2, LEVLFQGP), and a point mutant corresponding to the barcode (**Fig. S2D**). The peptide barcode formula was modified from previous work^27^ as follows: GX_4_WZ_4_R where G=glycine, X_4_=randomized region 1, W=tryptophan, Z_4_=randomized region 2, R=arginine (**Fig. S2A**). G and W contribute to solubility and support chromatographic separation, respectively. R provides charge for efficient LC-MS detection and allows for peptide formation following trypsinization. The randomized regions (X_4_ and Z_4_) serve to distinguish the identity of the assigned mutant and can contain A, N, D, E, Q, G, F, P, S, T, W, Y, V, I, or L. Disallowed amino acids include K, R, H, M, and C. Exclusion of K and R prevent barcode truncation during trypsinization. H was omitted to maintain charge state. M and C, sulfur-containing amino acids, were excluded to avoid changes in physiochemical properties (e.g., oxidation, di-sulfide bridge formation, carbamoylation, etc.). Barcode sequences and properties can be found in **Table S2**.

Peptide barcode masses (Da) and hydrophobicity values (SSRC^32^) were calculated on randomly generated peptides in R using the protViz package (https://CRAN.R-project.org/package=protViz). Barcodes were further filtered to include only masses between 1100–1500 Da.

A subset of peptide barcodes (synthesized by IDT) was fused to WT MS2 coat protein by PCR, and the barcoded WT MS2 coat protein pool was cloned and expressed as protein. Barcodes from the plasmid pool were sequenced by Illumina. Peptide barcodes were enriched by Ni-NTA agarose beads (Qiagen) as previously described^27^, and analyzed by LC-MS. Illumina sequencing counts and LC-MS abundances were fit to a linear model and residuals were calculated. Barcodes were excluded from the usable set if their residual was ≥ ± 2 or if their percentile rank was in the top 5% or bottom 10% of the data.

### Synthesis of RBR-scan pools

An N-terminal 6x-His tag was added to DNA templates by PCR using Q5 Hot Start polymerase (NEB). The His-modified template was PCR amplified with a deoxynucleotide mix containing dUTP in place of dTTP. Barcoded, N-terminal mutant fragments (F primer: 5’-XhoI site, peptide barcode DNA sequence, annealing region to 6x-His sequence-3’; R primer: point mutation) and their overlapping 3’ partners (F primer: point mutation complement; R primer: 5’-NcoI site, annealing region to the gene-3’) were synthesized by PCR with Q5U Hot Start DNA polymerase (NEB) using 15 pg template and 50 nM primers. For OE-PCRs with Kapa polymerase (Roche), complementary fragments were diluted 1:50 and combined with external primers. Thermolabile USER II (NEB) was added to the reactions for 60 min at 37 °C prior to PCR amplification. All OE-PCRs were pooled at equal volumes and gel purified. Purified pools were digested using XhoI and NcoI and ligated into a pMAL-c5P vector with T4 DNA ligase (Enzymatics) (insert:vector ratio for MS2 coat protein was 7:1 and 3:1 for SRSF2). Ligations were dialyzed using a 0.025 µm mixed cellulose ester membrane (MF Millipore VSWP01300) against deionized water. Electrocompetent NEB-10β *E. coli* (NEB C3020K) were transformed with ¼ of the dialyzed ligation reaction using the manufacturer’s protocol. Plasmids were extracted from an *E. coli* culture containing an average of 82,500 colony forming units, providing a 875x coverage for each individual mutant within a pool of 96 mutants. Plasmid midi preps were prepared using the manufacturer’s protocol (Macherey-Nagel). Plasmid quality was confirmed by PCR amplifying the plasmid backbone from the empty vector and the RBR-scan pools. Libraries for nanopore were amplified from the plasmid pool with Q5 Hot Start polymerase using the external primers from OE-PCR (**Table S4**).

### Recombinant protein expression and purification

Individual GST and MBP fusion proteins were expressed in BL21(DE3) for 16 hours at 16 °C. For GST proteins, cells were lysed in B-PER™ Complete Bacterial Protein Extraction Reagent (Thermo Fisher Scientific 89821), loaded onto a glutathione-sepharose gravity column, washed with PBS, and eluted in the presence of 10 mM glutathione. For MBP-MS2 coat proteins, cells were lysed in B-PER, loaded onto an amylose-sepharose gravity column, washed with column buffer (20 mM Tris, pH 7.5_RT_, 200 mM NaCl, 1 mM EDTA, 1 mM DTT), and eluted with 50 mM maltose in column buffer. Cells containing MBP-SRSF2 proteins were lysed in high salt column buffer (20 mM Tris, pH 8_RT_, 700 mM NaCl, 1 mM EDTA, 1 mM DTT) with sonication (Branson sonifier, 40% output, 8 x 15s on/1 min off), purified as described above, and eluted with high salt column buffer containing 50 mM maltose. Protein fractions were analyzed for purity by SDS-PAGE followed by Coomassie blue stain and pooled. Buffers were exchanged by dialysis or Amicon Ultra Centrifugal Filters (Millipore Sigma) against BC_100_ (20 mM Tris pH 7.9_4°C_, 100 mM KCl, 0.2 mM EDTA, 10% glycerol). Proteins were quantified by A280 and Bradford assays. Protein purity was confirmed by SDS-PAGE and Coomassie (see figures). Expression and purification conditions for RBR-scan pools were identical, except proteins were expressed in electrocompetent BL21(DE3).

### *In vitro* RNA binding assays

Biotinylated RNA was synthesized by *in vitro* transcription using a template containing three MS2 stem loop repeats (HiScribe T7 High Yield RNA Synthesis Kit, NEB E2040S) following manufacturer protocol except UTP was added at a ratio of 6.5:1 UTP:biotin-16-UTP (Sigma-Aldrich, 11388908910). RNA was purified using phenol chloroform extraction. RNA sequence information can be found in **Table S3**.

Prior to binding assays, RNA was refolded (2 minutes at 95°C, 2 minutes on ice, 20 minutes room temperature) in BTE buffer (10 mM bis-tris pH 6.7, 1 mM EDTA). Proteins were incubated with RNA in binding buffer (10 mM Tris pH 8_RT_, 200 mM KCl, 5 mM MgCl_2_, and 1 mM DTT) for 30 min on ice. Protein–RNA complexes were captured by incubating with streptavidin-conjugated dynabeads (Thermo Fisher) for 15 min at room temperature. Beads were washed three times with wash buffer (10 mM Tris pH 8_RT_, 200 mM KCl, 5 mM MgCl_2_, and 1 mM DTT, 0.05% IGEPAL CA-630) and three times with binding buffer.

### Fluorescence polarization assays

Fluorescence polarization (FP) assays were set up in 96-well black microplates (Greiner Bio-One) in a total reaction volume of 50 µL by titrating increasing concentrations of SRSF2 RRM proteins, ranging from 16 nM to 1 µM, into a buffer containing 10 mM HEPES pH 7.6, 100 mM KCl, 5 mM MgCl_2_, 2 mM DTT, 0.05% Tween-20, and 5 nM 5’ Alexa Fluor 488 labeled RNA probes, (IDT, **Table S3**), and incubated at 30°C for 30 min. FP readings were measured using a PerkinElmer Envision plate reader with excitation and emission wavelengths of 480 nm and 535 nm, respectively. Each experiment was performed in duplicate, and fluorescence values were normalized against the background fluorescence of the buffer containing the probe alone. Dissociation constants were calculated by fitting the polarization data to a one-site binding model using non-linear regression analysis in GraphPad Prism™ (version 10).

### Electrophoretic mobility shift assays (EMSAs)

RNA with a 5’ Alexa488 fluorophore was purchased from IDT (**Table S3**). Proteins were incubated with 50 nM RNA for 30 min at 30 °C in binding buffer (10 mM HEPES pH 7.6, 5 mM MgCl_2_, 100 mM KCl, 2 mM DTT, 50% glycerol, 0.05% Tween 20, 1g/L yeast tRNA (Thermo Fisher AM 7119)). Protein-RNA complexes were loaded onto 10% acrylamide mini gels pre-run at 4 °C for 30 min at 100 V in 1x TAE (40 mM Tris-acetate pH 7.8, 2.5 mM EDTA). Electrophoresis was performed at 4 °C for 3 min at 120 V and then for 1 hr at 100 V. Fluorescent RNA was imaged using an Amersham Imager600 (GE Healthcare).

### Mass Spectrometry

50 mM ammonium bicarbonate pH 8 was added to proteins on-bead or in buffer. Proteins were reduced (10 mM DTT, 55°C, 30 min), alkylated (25 mM iodoacetamide, 30 min in the dark), and trypsinized at 1:10 (w/w) enzyme:protein ratio overnight at 37°C (Sequencing Grade Modified Trypsin, Promega V511A). The pH was reduced to 2 with trifluoroacetic acid to stop digestion. Peptides were desalted, lyophilized, and resuspended in 4% acetonitrile containing 0.1% formic acid prior to LC-MS analysis.

Mass spectra for peptide barcodes were obtained with a Dionex Ultimate 3000 UHPLC system (Thermo Fisher Scientific) coupled to a Q-Exactive HF mass spectrometer (Thermo Fisher Scientific). Peptides were loaded onto an in-house silica capillary column (75 µm inner diameter) packed with C18 resin (ReproSil-Pur C18-AQ 2.4-μm resin; Dr. Maisch GmbH, Ammerbuch, Germany). Peptides were eluted with a gradient of 5%-45% B (60 min) followed by 95% B (10 min) and 95%-5% B (10 min) (mobile phase A: 0.1% formic acid, mobile phase B: 80% acetonitrile, 0.1% formic acid). Flow rate was 300 nL/min. Data were collected in data-dependent acquisition mode. Full-scan MS settings were as follows: scan range, 300–1,500 (m/z; mass-to-charge ratio); resolution, 120,000; AGC target 1E6; Maximum IT, 25 ms. MS2 settings were as follows: resolution, 15,000; AGC target 1E5; maximum IT, 100 ms; fragmentation was enforced by higher-energy collisional dissociation with normalized collision energy of 30; loop count, top 15; isolation window, 2.0 m/z; fixed first mass, 100 m/z; minimum AGC target, 800; charge exclusion: unassigned and 1; peptide match, preferred; exclude isotope, on; dynamic exclusion, 10 s.

Mass spectrometry data was analyzed using MetaMorpheus version 1.0.5 (https://github.com/smith-chem-wisc/MetaMorpheus) ^62^ using FASTA files containing peptide barcodes, MS2 coat protein or SRSF2, and common contaminants. The following search settings were used: protease = trypsin; search for truncated proteins and proteolysis products = false; maximum missed cleavages = 2; minimum peptide length = 7; maximum peptide length = 60; initiator methionine behavior = variable; fixed modifications = carbamidomethyl on C, carbamidomethyl on U; variable modifications = oxidation on M; max mods per peptide = 1; max modification isoforms = 1,024; precursor mass tolerance = ±10.0000 PPM; product mass tolerance = ±0.0200 absolute; report PSM ambiguity = true. Results were filtered for score (> 5) and FDR (1 %).

### Illumina sequencing

P5 and P7 adapter sequences encoding custom indexes were introduced by PCR amplification of the plasmid pool. The PCR product was size-selected by gel extraction (500 bp). Libraries were quantified by qPCR and sequenced on an Illumina NextSeq 500.

### Nanopore sequencing

Peptide-barcoded mutants were amplified from RBR-scan plasmid pools using Kapa HiFi Polymerase (Roche, 07958927001). Libraries for nanopore were prepared with the Native Barcoding Kit according to manufacturer’s instructions with minor adjustments (Oxford Nanopore Technologies, SQK-NBD114.24). 200 fmol of PCR product from RBR-scan pools was end-repaired using End Repair Enzyme Mix (Enzymatics, Y9140-LC-L), tailed with deoxyadenine using Klenow (Enzymatics, P7010-HC-L), and ligated to nanopore Native Barcodes. Nanopore libraries were pooled, size selected by SPRI (1.0x clean) and ligated to the nanopore sequencing adapter. Residual sequencing adapter was removed by SPRI. DNA was quantified by Qubit fluorimetry (Thermo Fisher) and 10–20 fmol were loaded onto the flow cell (Oxford Nanopore Technologies, FLO-MIN114). Sequencing rate = 400 bps.

### Nanopore data analysis

Bases were called using Dorado in mode dna_r10.4.1_e8.1_400bps_sup. Reads were subsequently filtered and aligned to the DNA sequence for MS2 coat protein or SRSF2. Mutation frequencies were calculated as the percentage of reads containing the correct mutant codon out of the total reads associated with each barcode. Reads were further categorized as the desired mutation, WT, or other (i.e., neither the expected mutant nor the WT sequence). Mutants linked to barcodes with ≤ 10 reads or that contained ≥ 50% of reads sourced from WT were excluded from mass spectrometry analysis. In some cases, barcodes associated with mutations outside their expected pool were reassigned to the mutation they were linked to. Specifically, 11 out of 192 barcodes were mismatched for MS2 coat protein, and 4 out of 68 were mismatched for SRSF2.

### LC-MS data analysis

Barcodes not detected by nanopore sequencing were excluded from LC-MS analysis. Barcode abundances were normalized to total abundance per run to minimize within-sample variability. Barcode depletion (log_2_-fold change) was calculated by normalizing the mean pull-down:input abundance ratio to the mean WT barcode abundance.

## Data accessibility

The mass spectrometry proteomics data have been deposited to the ProteomeXchange Consortium via the PRIDE^63^ partner repository with the dataset identifier PXD062465. Nanopore sequencing data have been deposited to the GEO repository with the accession number GSE293492.

## Author contributions

J.A.T, R.B. and B.A.G. developed the project. J.A.T and R.B. designed all experiments. J.A.T. performed all experiments except for nanopore and fluorescence polarization which were conducted by J.F.D. and S.D.M., respectively. E.J.S. provided code and data analysis related to sequencing. K.S. provided input on FP and EMSA experiments. L.N.R. performed cell culture experiments. B.A.G. provided mass spectrometry supervision. J.A.T. and R.B. wrote the manuscript with input from all other authors.

## ACKNOWLEDGMENTS

The authors thank A. Scacchetti, K. Lynch, and M. Tracey for technical support. J.A.T. was supported by the NIH F31 HD111270, J.F.D. was supported by NIH training grant T32 HD083185, and L.N.R. was supported by NSF training grant DGE 2236662. R.B. acknowledges support by the NIH (R35 GM153281). K.S. acknowledges support by the NIH (R01 GM143229). B.A.G. acknowledges support by the NIH (R01 HD106051 and R01 AI118891).

## SUPPLEMENTAL FIGURE LEGENDS

**Supplemental figure 1.**
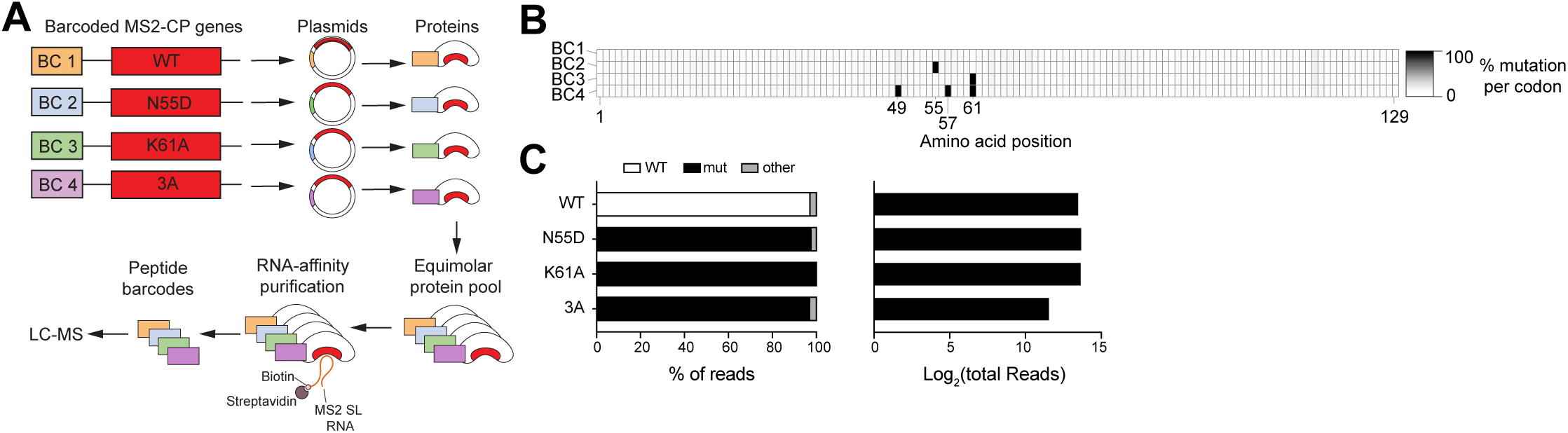
RBR-scan with individual MS2 coat proteins. (A) Experimental scheme of low-throughput RBR-scan proof-of-concept experiment. BC, barcode. 3A, R49A-K57A-K61A mutant. (B) Heatmaps of mutation frequencies (% of reads) at each amino acid (x-axis) for plasmid pool of commercially synthesized MS2 coat protein variants paired to different barcodes (y-axis), as determined by nanopore sequencing. Positions of point mutations are indicated. (C) Frequency of mutant reads associated with each barcodes. Desired mutation (mut, black) is the correctly assigned mutation, WT sequences are in white, and other reads (gray) indicate mutation that do not correspond to either.

**Supplemental figure 2.**
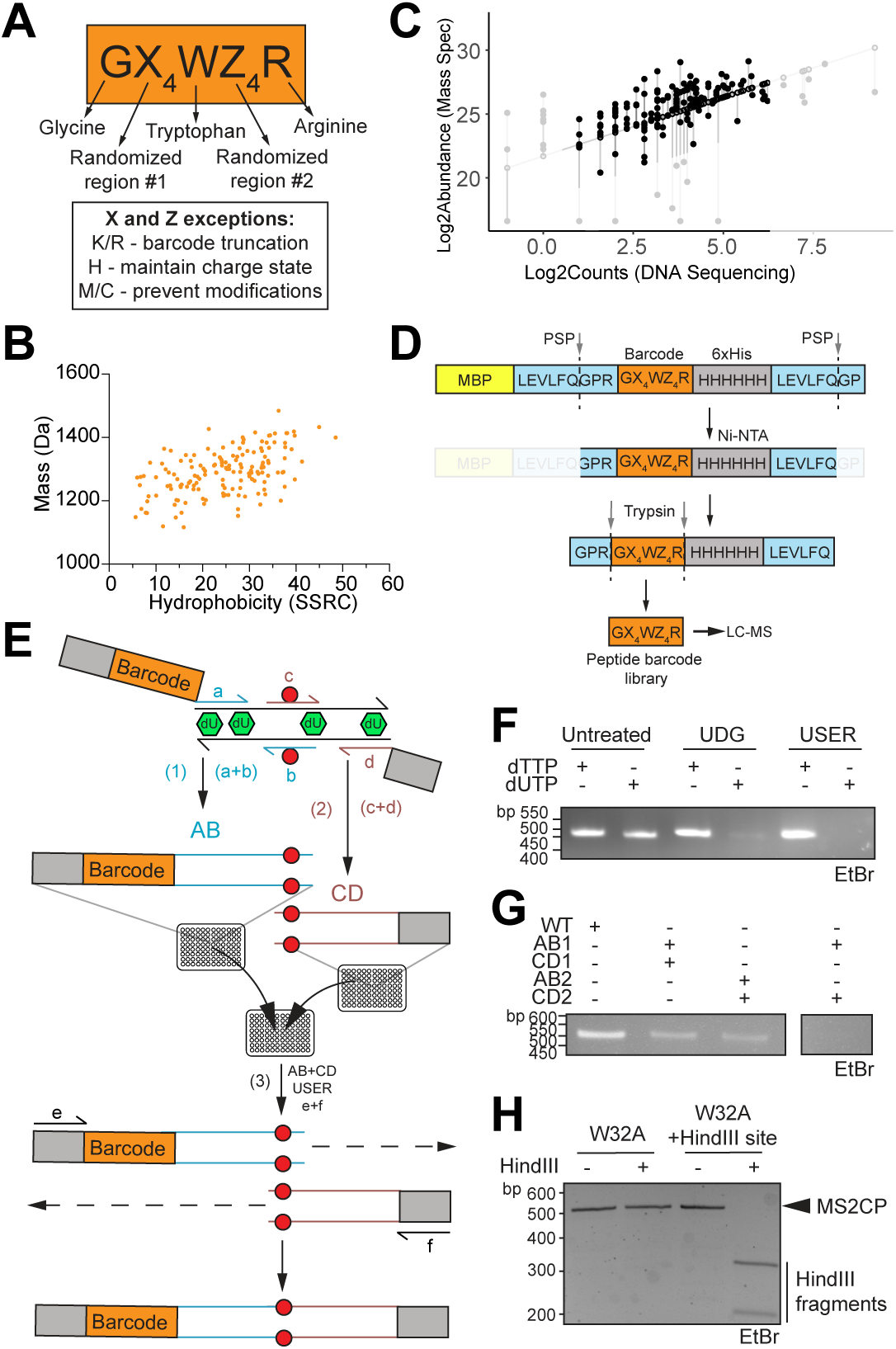
Barcode and assay design for high-throughput scanning. (A) Peptide barcode design. See methods. (B) Hydrophobicity (calculated by sequence-specific retention calculator (SSRC)^32^) plotted against peptide barcode mass (Da) for 155 peptide barcodes designed *in silico*. (C) Peptide barcode analysis of barcodes attached to WT MS2 coat protein. Comparison of the log_2_(counts) from Illumina sequencing (x-axis) and log_2_(abundances) from mass spectrometry (y-axis) of 155 peptide barcodes. Horizontal lines indicate residual values. Barcodes were excluded from RBR-scan experiments (faded points) if they were outside of the residual cutoff (± 2). (D) MBP-fusion protein pools were designed to contain N-terminal peptide barcodes flanked by two PreScission Protease (PSP) sites and a 6xHis-tag. Barcodes of PSP-digested pro-peptides enriched by Ni-NTA were trypsinized to yield peptide barcodes. See methods and **Fig. S2A** for barcode composition rules. (E) High throughput OE-PCR scheme. Involves three PCRs where PCRs 1 and 2 use a WT template and external primers a and d paired with internal mutant primers b and c to generate fragments AB and CD, respectively. Primer a contains the peptide barcode DNA sequence and is different for each mutant. The WT template used in fragment PCRs 1 and 2 is made with dUTP (green) so that it can be enzymatically degraded by uracil-specific excision reagent (USER) in PCR 3. PCR 3 incorporates external primers e and f instead of a and d as an additional safeguard against contamination from the WT template. All PCRs were conducted in multi-well plates to maintain high throughput. Red circle indications mutation. (F) Digestion of MS2 coat protein template with uracil-DNA glycosylase (UDG) or USER. (G) OE-PCR with mismatched AB and CD fragments. Combination of fragments AB1 with CD1 and AB2 with CD2 form PCR products at the expected size (500 bp), whereas the combination of fragments AB1 with CD2 does not. (H) Restriction digest assay for OE-PCR products from W32A with and without HindIII site (expected size of full length MS2 CP: 500 bp, expected sizes of HindIII fragments: 193 bp and 307).

**Supplemental figure 3.**
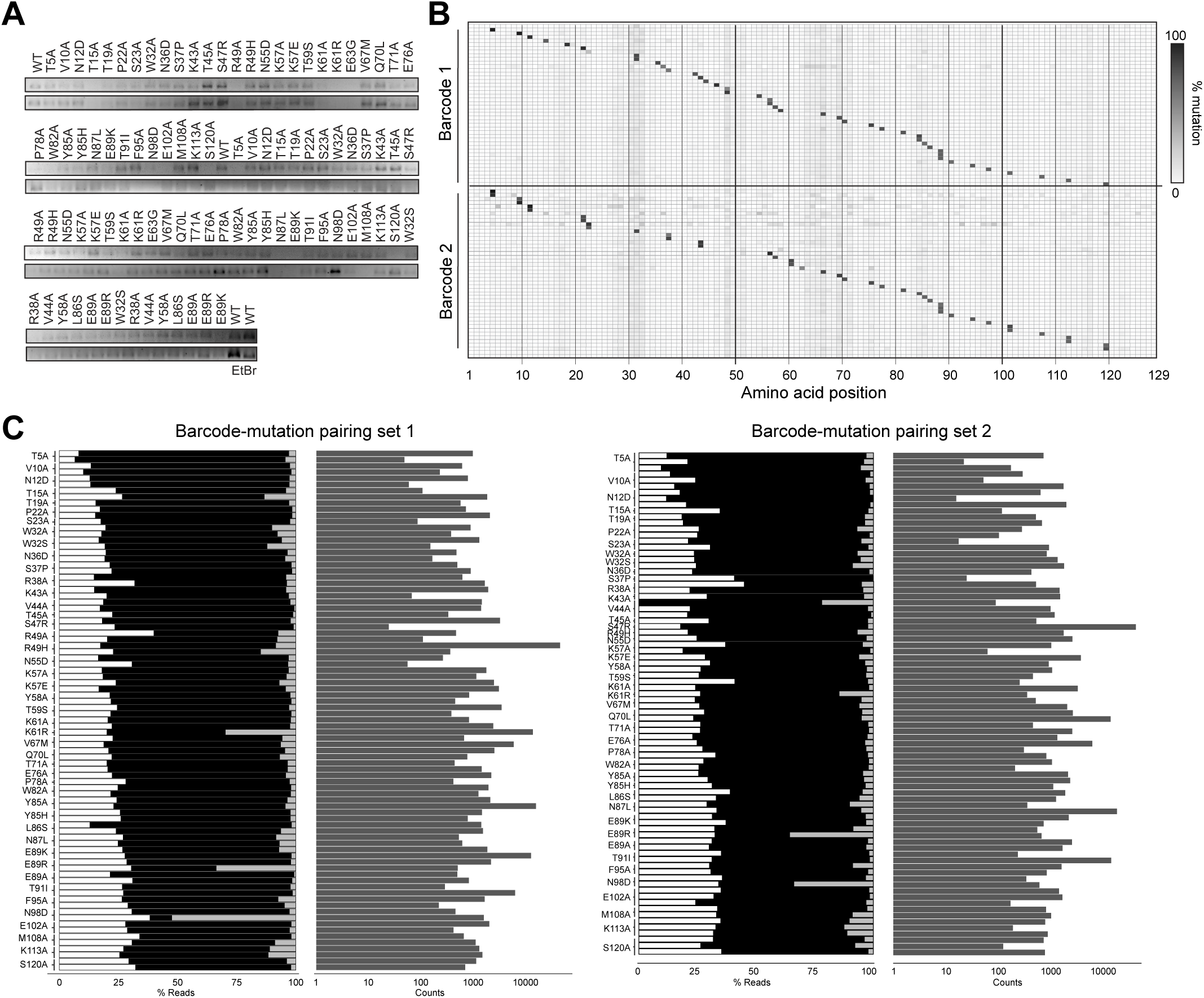
Verification of batch cloning and protein pools. (A) MS2 coat protein OE-PCR products from two different barcode-mutation configurations. (B) Alternate barcode-mutation configuration for MS2 coat protein RBR-scan pool (see Fig. 2C) Mutation frequencies (% of reads) of each amino acid (x-axis) along the MS2 coat protein sequence. Barcoded MS2 coat protein variants are shown (y-axis), and barcodes 1 and 2 denote two different peptide barcodes and codons for each amino acid within the same pool. (C) Source of nanopore reads associated with each peptide barcode (left) as well as the number of reads per barcode (right). Read source categories are desired mutation (mut, black), WT (white), and other (grey). Mut, mutant.

**Supplemental figure 4.**
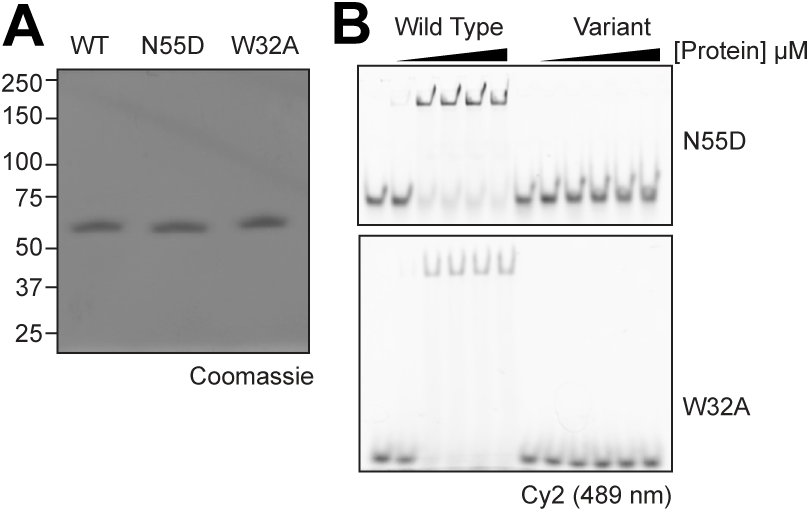
RBR-scan new mutant validation. (A) Coomassie stain of individual MBP-fused MS2 coat proteins (predicted molecular weight: 60.5 kDa). (B) EMSAs with titrations of recombinant MS2 coat proteins and 50 nmol Alexa488-MS2 stem loop RNA in the presence of 1 g/L yeast tRNA. Complexes were separated on native gels, and RNA was imaged by Cy2 excitation. Data is representative of ≥2 experiments. Replicate for Fig. 3C.

**Supplemental figure 5.**
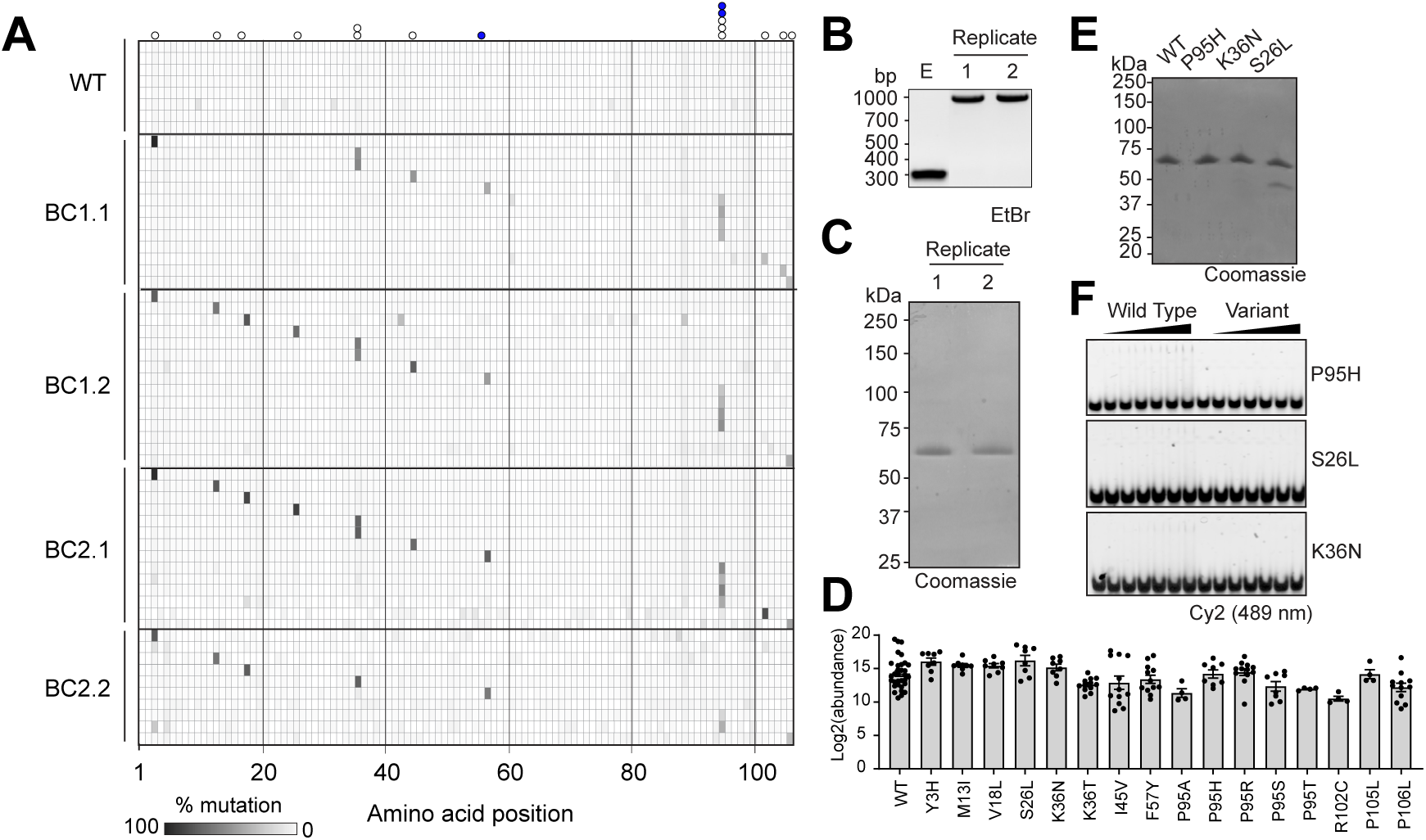
Batch cloning and protein expression of SRSF2 RBR-scan pools. (A) Mutation frequencies (% of reads) of each amino acid (x-axis) along the SRSF2 sequence. Barcoded MS2 coat protein variants are shown (y-axis), where WT barcodes as well as barcode 1 from barcode-mutation pair set #1 (BC1.1), BC1.2, BC2.1, and BC2.2 are indicated. Barcodes within a set (i.e., barcode 1 and barcode 2) are associated with different codons for the same amino acid-coding mutation. Sets refer to different barcode-mutation pairings. Positions of intended mutations are indicated at the top of the heatmap (white circles, unknown phenotype; blue circles, known RNA-binding mutant). (B) PCR products from two RBR-scan plasmid libraries (expected size: 1027 bp) and an empty vector (expected size: 278). E, empty vector. (C) Coomassie stain of MBP-fused peptide barcoded SRSF2 pools (predicted MW: 72 kDa). (D) Mass spectrometry analysis of peptide barcode abundances SRSF2 pools. Point mutations indicated on x-axis. Bars represent the mean log_2_(barcode abundance). Dots represent abundances from individual barcodes across replicates. Data from 8 biological replicates. (E) Coomassie stain of individual MBP-fused SRSF2 variants. (F) EMSAs with titrations of recombinant SRSF2 RRMs and 50 nmol Alexa488-MELK RNA in the presence of 1g/L yeast tRNA. Complexes were separated on native gels, and RNA was imaged by Cy2 excitation. Data is representative of ≥2 experiments. Replicate for Fig. 4D.

## SUPPLEMENTAL TABLE LEGENDS

**Table S1. Published dissociation constants for MS2CP**

**Table S2. Barcode-mutation assignments**

**Table S3. Barcode amino acid sequences and properties**

**Table S4. Primers and oligos**

